# Deleterious variation mimics signatures of genomic incompatibility and adaptive introgression

**DOI:** 10.1101/221705

**Authors:** Bernard Y. Kim, Christian D. Huber, Kirk E. Lohmueller

## Abstract

While it is appreciated that population size changes can impact patterns of deleterious variation in natural populations, less attention has been paid to how population admixture affects the dynamics of deleterious variation. Here we use population genetic simulations to examine how admixture impacts deleterious variation under a variety of demographic scenarios, dominance coefficients, and recombination rates. Our results show that gene flow between populations can temporarily reduce the genetic load of smaller populations, especially if deleterious mutations are recessive. Additionally, when fitness effects of new mutations are recessive, between-population differences in the sites at which deleterious variants exist creates heterosis in hybrid individuals. This can lead to an increase in introgressed ancestry, particularly when recombination rates are low. Under certain scenarios, introgressed ancestry can increase from an initial frequency of 5% to 30-75% and fix at many loci, even in the absence of beneficial mutations. Further, deleterious variation and admixture can generate correlations between the frequency of introgressed ancestry and recombination rate or exon density, even in the absence of other types of selection. The direction of these correlations is determined by the specific demography and whether mutations are additive or recessive. Therefore, it is essential that null models include both demography and deleterious variation before invoking reproductive incompatibilities or adaptive introgression to explain unusual patterns of genetic variation.

## Introduction

There is tremendous interest in quantifying the effects that demographic history has had on the patterns and dynamics of deleterious variation and genetic load (Kirkpatrick and Jarne 2000; Gazave et al. 2013; Lohmueller 2014a; Lohmueller 2014b; Henn et al. 2015; Brandvain and Wright 2016; Simons and Sella 2016). Several studies have suggested that recent human demography has had little impact on load (Simons et al. 2014; Do et al. 2015) while others have suggested weak, but subtle differences between human populations (Casals et al. 2013; Fu et al. 2014; Gravel 2016; Henn et al. 2016; Pedersen et al. 2017). All of these studies have typically focused on how population size changes, such as expansions and bottlenecks, have affected deleterious variation. Other types of complex demography, however, have received considerably less attention.

In particular, gene flow may be important for shaping patterns of deleterious variation. Population admixture, or hybridization between closely related species, appears to be quite common in nature (Payseur and Rieseberg 2016) and has a significant role in shaping human genomes (Wall and Yoshihara Caldeira Brandt 2016). Gene flow alone can subtly change the effects of selection on deleterious variation (Gravel 2016), but should have notable fitness consequences if deleterious variation is distributed differently between admixing populations. For example, Neanderthals likely had a higher genetic load than coincident human populations due to the former’s smaller long-term population size (Do et al. 2015; Harris and Nielsen 2016; Juric et al. 2016). As a result, gene flow from Neanderthals into the ancestors of modern humans could have increased the genetic load of some human populations by 0.5% (Harris and Nielsen 2016), and selection should have removed deleterious Neanderthal ancestry over time. In contrast, domesticated species likely have increased genetic load due to domestication bottlenecks and hitchhiking with artificially selected variants (Marsden et al. 2016; Liu et al. 2017; Moyers et al. 2017). Then, gene flow from their wild counterparts should alleviate the genetic load of domesticated species and increased levels of introgression should be observed (e.g. Wang L, Beissinger TM, Lorant A, Ross-Ibarra C, Ross-Ibarra J, Hufford M, unpublished data, https://www.biorxiv.org/content/early/2017/03/07/114579, last accessed Nov. 13, 2017).

If effects of gene flow can be modulated by the consequences of deleterious variation, selection on deleterious variants may provide an alternative explanation for patterns of introgression that are usually attributed to processes such as the evolution of reproductive incompatibility or adaptive introgression. Patterns such as the depletion of introgressed ancestry in regions of low recombination (Sankararaman et al. 2014) might be instead partially explained by purifying selection removing deleterious variation and partially by reproductive incompatibilities (Sankararaman et al. 2014; Harris and Nielsen 2016; Juric et al. 2016; Vernot et al. 2016). Rampant adaptive introgression may create the opposite pattern where there is increased amounts of introgressed ancestry in regions of low recombination (e.g. Pool 2015; Corbett-Detig and Nielsen 2017). Alternatively, this pattern could be due to selection favoring introgressed haplotypes from a larger population that carries fewer deleterious variants. The general extent of these effects in nature and their magnitude given realistic parameter combinations remains to be studied.

Another significant issue with many models incorporating deleterious variation is the assumption that fitness effects are strictly additive. A considerable proportion of strongly deleterious new mutations are likely to be fully recessive or partially recessive (Simmons and Crow 1977; Agrawal and Whitlock 2011; Huber CD, Durvasula A, Hancock AM, Lohmueller KE, unpublished data, https://www.biorxiv.org/content/early/2017/08/31/182865, last accessed Nov. 13, 2017). In addition, if some proportion of deleterious recessive variants is private to a population, admixed populations should experience heterosis since deleterious variants are more likely to be found in a heterozygous state (Crow 1948). As a result, heterosis can increase effective migration rates between populations (Ingvarsson and Whitlock 2000) and may play a significant role in determining the fitness of highly structured populations (Whitlock et al. 2000). Heterosis may also increase introgression and the probability that introgressed ancestry will persist in an admixed population, even if the introgressed ancestry contains more deleterious variation (Harris and Nielsen 2016). The extent to which heterosis contributes to increases in the frequency of introgressed ancestry and confounds inference of adaptive introgression is also not well understood.

The objective of this study is to develop a clearer picture of the effect of deleterious variants on fitness and the dynamics of introgression in admixing populations. Further, we aim to understand how inferences of selection on introgressed ancestry are impacted by deleterious variation. Previous simulation and empirical work have shown that for at least some systems, deleterious variation is a significant factor modulating gene flow (Harris and Nielsen 2016; Juric et al. 2016; Wang et al. 2017), but few studies have investigated these questions outside of demographic models specific to a species or system. Our present study fills this void by presenting a series of simulations utilizing demographic models that generalize biological scenarios of interest. We include a realistic distribution of fitness effects for both additive and recessive mutation models. In addition, we examine how the relationship between introgressed ancestry and recombination rates, or functional content, is determined by the underlying demography.

## Results

### Forward simulations

We used SLiM 4.2.2 (Haller and Messer 2017) to simulate admixture in the presence of deleterious variation. In general, the underlying demographic model was a population split model with an ancestral population size (*N_a_*) of 10,000 diploids, where a single pulse of admixture occurs at an initial proportion of 5%, in one direction and for one generation, at some time after the split (**Figure 1, Table 1**). Throughout, we will refer to the subpopulation from which gene flow occurs as the *source* subpopulation, and the subpopulation which receives gene flow as the *recipient* subpopulation. Furthermore, we will refer to ancestry in the recipient subpopulation that originated in the source subpopulation as *introgression-derived ancestry*. We tracked the frequency of introgression-derived ancestry in simulations by placing marker mutations in one subpopulation, and use *p_I_* to denote the estimated total proportion of ancestry in the recipient subpopulation that is introgression-derived. The sizes of the subpopulations were varied as described in the forthcoming sections and in **Figure 1** and **Table 1**.

**Figure 1.**
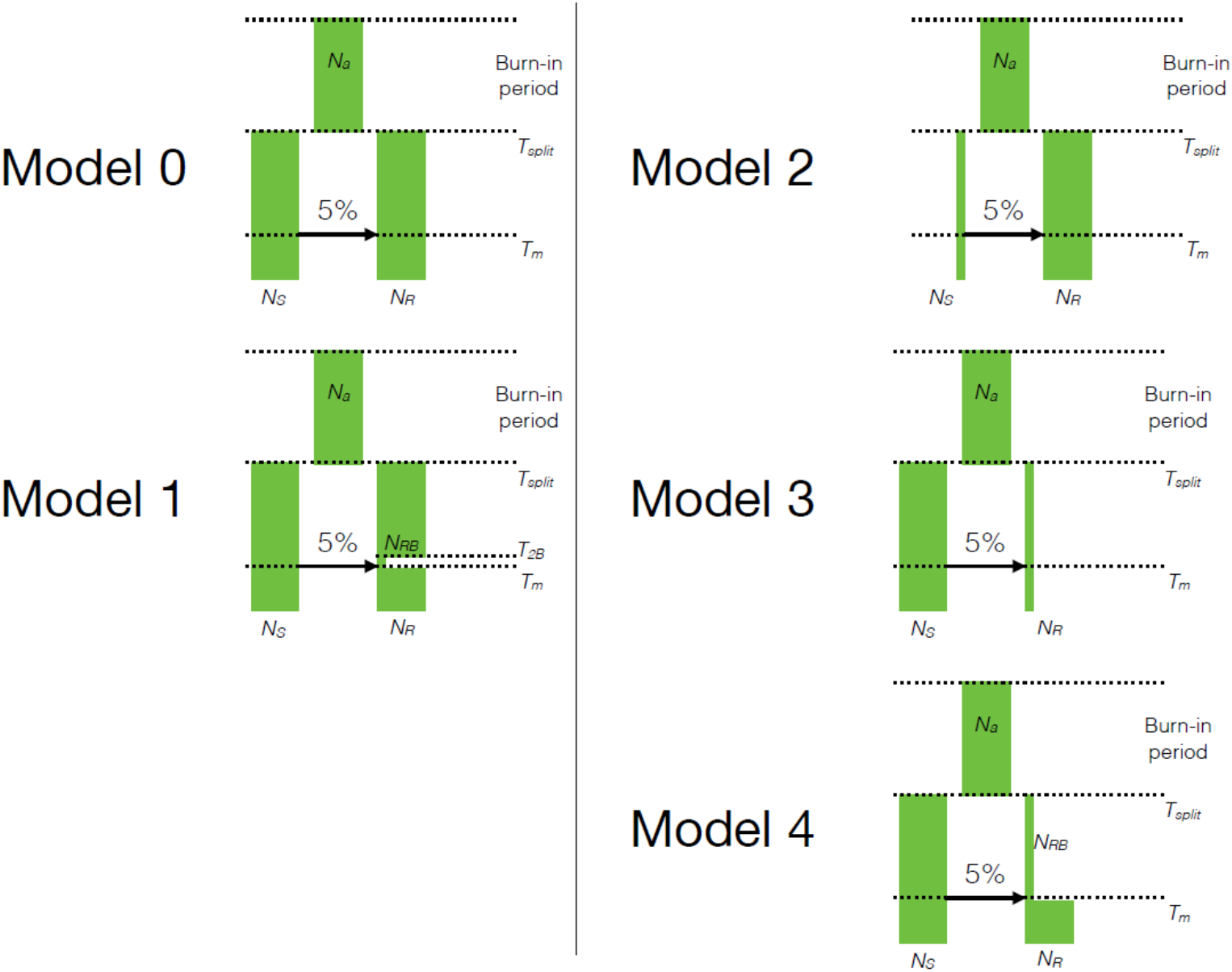
The demographic models used for the simulations. After a burn-in period of 100,000 generations, a single population diverged into two subpopulations. The demography of the subpopulations was modified in ways that changed the distribution of deleterious variation. 2,000 generations after the split, a single pulse of admixture occurred such that 5% of the ancestry of the recipient population came from the donor population. Arrows in each panel denote the direction of gene flow. The simulation was run for 10,000 additional generations after admixture. Population sizes were changed as shown for each model. See **Table 1** for specific parameter values for each model.

**Table 1.** Demographic parameters of the simulated models shown in Figure 1.

| Model | $N_a$ | $N_S$ | $N_{RB}$ | $N_R$ | $T_m$ | $T_{RB}$ | $T_{split}$ |
| --- | --- | --- | --- | --- | --- | --- | --- |
| Model 0 | 10,000 | 10,000 | -- | 10,000 | 1,000 | -- | 3,000 |
| Model 1 | 10,000 | 10,000 | 1,000 | 10,000 | 1,000 | 1,050 | 3,000 |
| Model 2 | 10,000 | 1,000 | -- | 10,000 | 1,000 | -- | 3,000 |
| Model 3 | 10,000 | 10,000 | -- | 1,000 | 1,000 | -- | 3,000 |
| Model 4 | 10,000 | 10,000 | 1,000 | 10,000 | 1,000 | -- | 3,000 |

NOTE.--All population sizes are in diploids and all times in generations from the present.
Parameters are defined as follows. $N_a$ : ancestral population size, $N_S$ : size of the source subpopulation, $N_{RB}$ : size of the bottleneck in the recipient population, $N_R$ : size of the recipient population, $T_m$ : time of migration, $T_{RB}$ : time at which the bottleneck in the recipient population began, $T_{split}$ : time at which the subpopulations diverged

Unless specified otherwise, we simulated approximately 5 Mb of sequence, with randomly generated genic structure (see Methods). The mutation rate (*μ*) was set at a constant rate of 1.5×10^−8^ per bp per generation. All simulated mutations were either neutral or deleterious, and only nonsynonymous mutations had nonzero selection coefficients. Deleterious mutations had selection coefficients (*s*) were all drawn from the same distribution of fitness effects (Kim et al. 2017). In other words, the selection coefficients did not depend on the population in which mutations occurred. No positively selected mutations were simulated in any of our models. We simulated additive (*h*=0.5) and recessive (*h*=0.0) mutations separately. Additive fitness effects were computed by multiplicatively calculating fitness at each locus. Recessive fitness effects were computed additively, but only at homozygous loci. All fitness effects were computed multiplicatively across loci. To assess the effect of recombination rate, we also varied the per-base pair per chromosome recombination rate, *r*, between sets of simulations (*r*∈{10^−6^,10^−7^,10^−8^,10^−9^}). See **Methods** for additional details on the simulations and calculation of fitness.

In each simulation, we recorded the fitness of each subpopulation, the proportion of ancestry that is introgression-derived, and other measures of genetic load (**Figures S1-S3**) at different time points. Differences in subpopulation fitness are presented as (*w_R_*/*w_S_*), which represents the relative differences in load between the recipient and source subpopulation. Subpopulation fitnesses are presented separately in **Figure S1**.

### The effect of admixture on deleterious variation

Our first step was to look at only the effects of gene flow on deleterious variation when the two subpopulations had identical sizes. Given that the populations were of identical size, they both should contain similar amounts of deleterious variation and similar genetic loads. Here, we simulated under a population split model with a size of 10,000 diploids in the ancestral and both divergent subpopulations (Model 0, **Figure 1**), where 20,000 (2*N_A_*) generations after the population split a single pulse of admixture occurred.

Under the additive fitness model, the fitness of the recipient subpopulation does not change due to gene flow (Model 0, **Figure 2**). At the time point of admixture, the two subpopulations have similar fitnesses due to identical population sizes (*w_S_* ≈ *w_R_*, **Figures 2** and **S1**), although many deleterious variants will be private to each subpopulation. In addition, gene flow does not change the mean number of deleterious variants per haplotype, since any introgression-derived haplotypes carry, on average, the same number of derived variants as background haplotypes (**Figure S2**).

**Figure 2.**
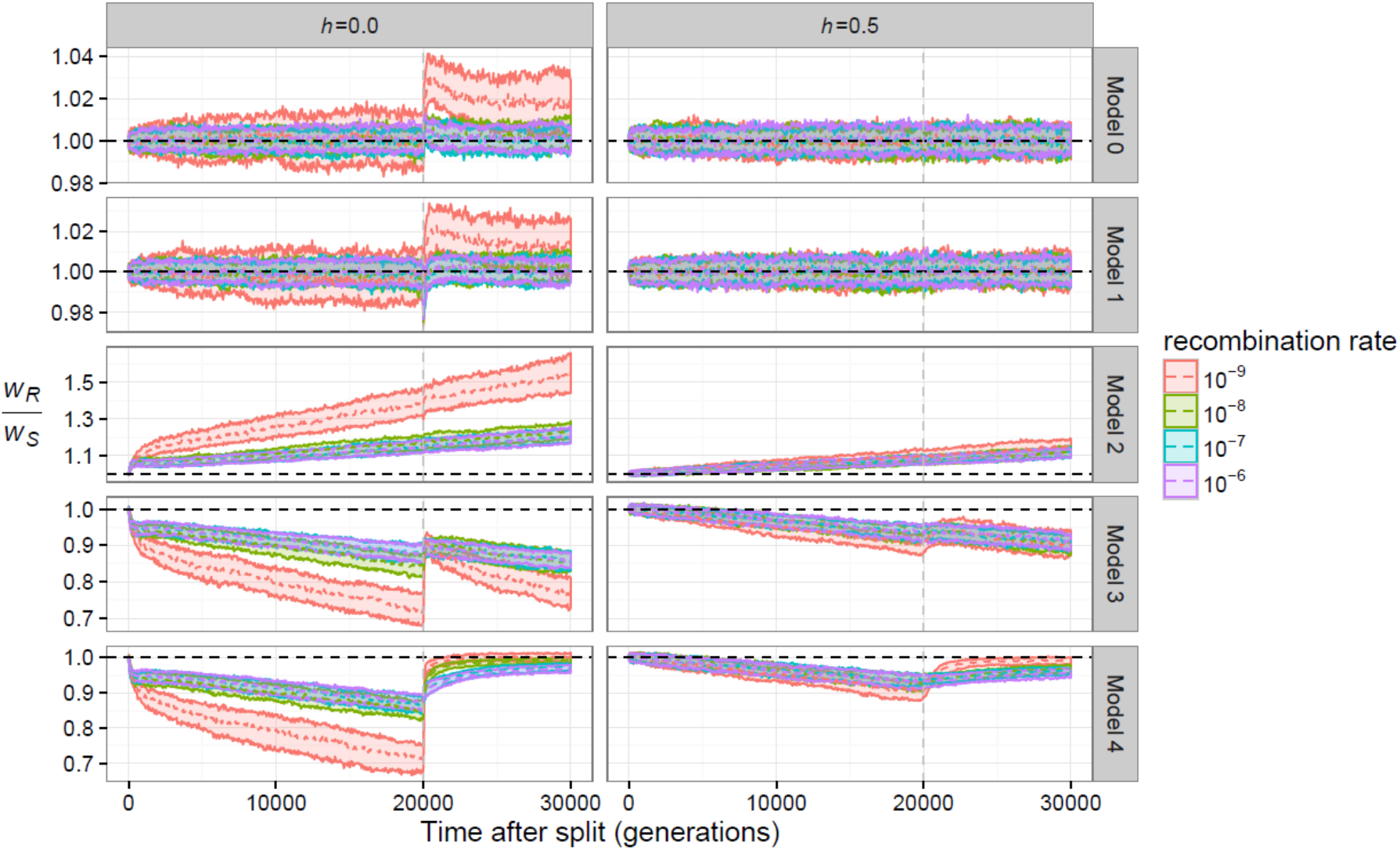
The change in mean fitness over time due to demography. Each individual plot depicts the ratio of the relative mean fitness of the recipient population (*w_R_*) to the source population (*w_S_*) for the demographic models shown in **Figure 1**. The median (dotted line) and the 25^th^ to 75^th^ percent quantiles are shown for 200 simulation replicates. The vertical grey line depicts the time of gene flow, and the horizontal black line depicts *w_R_*/*w_S_*=1. Different colors denote distinct recombination rates used in the simulations. Left panel denotes recessive mutations (*h*=0) while the right panel shows additive mutations (*h*=0.5).

When deleterious mutations are recessive instead of additive, admixture is predicted to generate heterosis in hybrid individuals, particularly when some proportion of segregating variants is private to each subpopulation (Crow 1948; Whitlock et al. 2000; Harris and Nielsen 2016). Indeed, 20,000 generations after the population split, many of the deleterious variants in our simulations should be private to one subpopulation (**Figure S4**). At the time of gene flow, both subpopulations have declined in fitness to a similar degree (*w_R_* ≈ *w_S_*, Model 0, **Figure 2**). After admixture, homozygosity in the recipient population is immediately decreased (**Figure S3**), with a corresponding increase in the fitness of the recipient population (**Figures 2** and **S1**). Importantly, the number of derived deleterious variants per haplotype in the recipient subpopulation is unaffected by gene flow (**Figure S2**). After the initial increase in fitness due to admixture, the decline in fitness following admixture is slow, such that fitness will be greater than the pre-admixture value for many generations following admixture (Model 0, *h*=0, **Figure S1**). Notably, the fitness increase conferred by heterosis is the most pronounced (≈2.5% increase) and lasts the longest (>10,000 generations after admixture) in simulations with low amounts of recombination (*r*=10^−9^). This occurs for two reasons. First, fitness declines more quickly when recombination rates are low (Muller 1964), resulting in stronger selection against non-admixed individuals. Second, introgression-derived haplotypes remain intact, maximizing the heterotic advantage of admixed individuals.

The effect of selection on deleterious variation also influences the fraction of ancestry that is introgressed in the recipient population. Considering that immediately following admixture, 5% of the ancestry in the recipient population is introgression-derived (*p_I_* =5%), it follows that in the absence of allele frequency changes due to selection the expected proportion of introgression-derived ancestry should remain E[*p_I_*]=5%. Our simulations show that this is the case when fitness effects are additive and both subpopulations have similar genetic loads (Model 0 in **Figure 3**). Therefore, selection does not favor haplotypes of a particular ancestry, and the long-term *p_I_* ≈ 5%.

**Figure 3.**
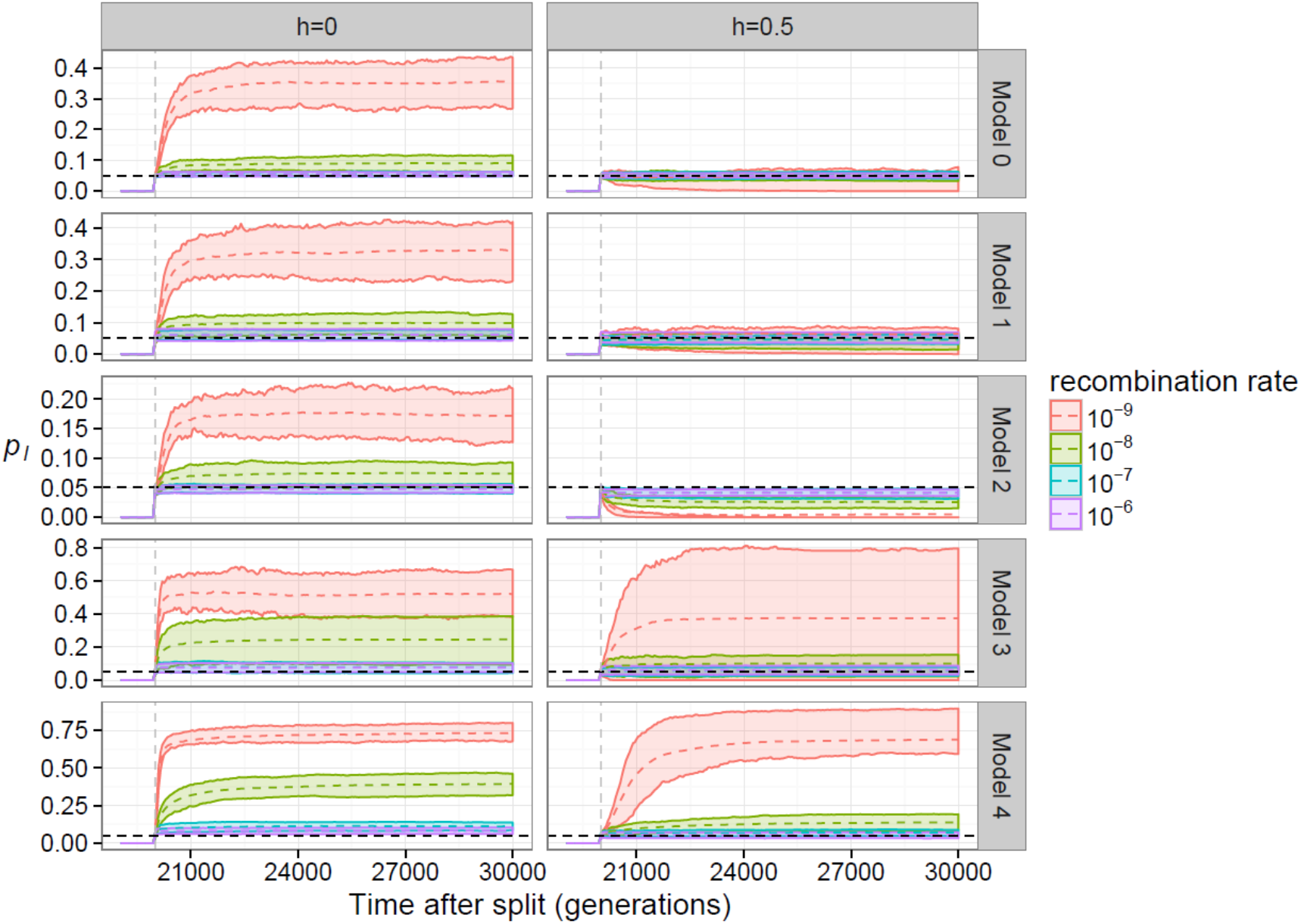
The frequency of introgression-derived ancestry (*p_I_*) in each model. Earlier generations are not shown since *p_I_*=0 prior to admixture. The mean (dotted line) and the 25^th^ to 75^th^ percent quantiles are shown for 200 simulation replicates. The vertical red line depicts the time of gene flow, and the horizontal black line depicts the initial admixture proportion of 0.05. Different colors denote distinct recombination rates used in the simulations. Left panel denotes recessive mutations (*h*=0) while the right panel shows additive mutations (*h*=0.5).

If deleterious mutations are recessive, introgression-derived ancestry increases in frequency as protective haplotypes rise in frequency through heterosis (*p_I_* > 5%; Model 0, *h*=0, **Figure 3**). The increase in *p_I_* is inversely related to the recombination rate. Specifically, the increase in *p_I_* is greatest (average *p_I_*~35% at 10,000 generations after the split, **Figure 3**) for the lowest recombination rate *r*=10^−9^, (**Figures 2** and **S1**). This effect is still observed even when the simulated recombination rate is greater than 10^−9^, but the magnitude of the effect is far less pronounced, with increases to *p_I_* ≈ 6-9%. These results also corroborate studies which showed that heterosis can increase effective migration rates (Ingvarsson and Whitlock 2000) or enhance the introgression of linked deleterious variants (Harris and Nielsen 2016). However, our results additionally show that heterosis should be greater when recombination rates are low.

### Short-term bottlenecks have little influence on the dynamics of introgression

Next, we investigated how short-term bottlenecks might affect fitness and patterns of introgression by adding a short bottleneck into one subpopulation of the split model (Model 1, **Figure 1**). Specifically, we added a bottleneck of size 1,000 diploids, or a 10-fold reduction in population size, in the recipient subpopulation for the 50 generations immediately preceding the admixture event. All population sizes remained at 10,000 diploids otherwise.

In this model, the additive genetic load was insensitive to the short bottleneck (Model 1, **Figures 2** and **S1**). Although some proportion of the deleterious variants is lost in the bottleneck, the average number of deleterious variants per haplotype is unchanged (Lohmueller 2014b; Simons et al. 2014; Simons and Sella 2016), and the additive load in each population is the same (*w_S_* ≈ *w_R_*, **Figures 2** and **S1**). Therefore, gene flow has no effect on the additive load or the number of deleterious variants per haplotype (**Figure S2**) in the recipient population, and there is no selection on introgressed ancestry in this model.

We found broadly similar patterns when deleterious mutations were recessive, with some important differences. Our simulations show that the bottleneck causes an additional ~2% decline in the recipient population’s mean fitness prior to admixture (Model 1, **Figure S1**) due to a small increase in homozygosity (**Figure S3**). However, the average number of deleterious variants per haplotype is unaffected by any selection against the increased proportion of homozygotes (Model 1, **Figure S2**). Fitness increases quickly after admixture, but consistently remains slightly less than the model without a bottleneck, suggesting that any change in load due to an increased number of homozygous sites is mostly cancelled out by the increased heterozygosity that results from admixture (**Figure S3**). The magnitude of the fitness increase from admixture is again inversely related to the recombination rate, in a manner similar to that in the model without a bottleneck (Model 0).

The frequency of introgression-derived ancestry was largely unaffected by the short bottleneck. When fitness effects were additive, the average frequency of introgressed ancestry remained at the initial admixture proportion of *p_I_* = 5%, 10,000 generations after the admixture event (Model 1, **Figure 3**). When fitness effects were recessive, introgression-derived ancestry increased in frequency by carrying protective alleles, similar to the simulations with identical subpopulation size. The same inverse relationship to the recombination rate was also observed. In the case of recessive mutations, the long-term frequency of introgression-derived ancestry (e.g. *p_I_* ≈ 33% for *r*=10^−9^, Model 1 in **Figure 3**) was similar but slightly lower than the model without a bottleneck (e.g. *p_I_* ≈ 35% for *r*=10^−9^, Model 0 in **Figure 3**).

### Long-term population contractions greatly influence the dynamics of introgression

At equilibrium, smaller populations will have a greater reductions of fitness due to deleterious variation than larger populations (Kimura et al. 1963). Therefore, a long-term reduction in population size should have different implications for fitness and the fate of introgression-derived ancestry than the short bottleneck described above.

To investigate the effect of long-term differences in population size and subsequent gene flow on patterns of deleterious alleles, we simulated a split model with the addition of a long-term reduction in population size. Immediately following the split, the size of one subpopulation was reduced 10-fold to 1,000 diploids (Models 2 and 3, **Figure 1**). After 20,000 additional generations, gene flow occurred in a single generation at an admixture proportion of 5%. In Model 2, the direction of admixture is from the small into the large population, and in Model 3 the direction of admixture is from the large into the small population.

As a consequence of long-term differences in population size, the additive fitness of the small subpopulation is 7-10% less than that of the large subpopulation (Models 2 and 3 in **Figures 2** and **S1**) at the time of admixture. In the additive fitness model, gene flow from the small to the large subpopulation (Model 2) resulted in a small fitness decrease (<1%) in the recipient subpopulation’s fitness, but had little effect on the average number of derived deleterious variants per haplotype (**Figure S2**). Gene flow from the large to the small subpopulation (Model 3) resulted in small increases (<1%) in fitness in the recipient subpopulation’s fitness, except (~5% increase) when recombination was low (**Figure S1**). In this case, selection for less deleterious haplotypes resulted in an overall decrease of the average number of deleterious variants per haplotype, but because the recipient population remained at a small size after admixture, load continued to accumulate afterward at the same rate (**Figures 2 and S1**).

When deleterious mutations were recessive, the effect of admixture on the recipient population’s fitness was determined both by differences in genetic load between populations and heterosis from admixture. Immediately prior to admixture, the recipient population’s fitness was approximately 10-30% greater (in Model 2) or less (in Model 3) than the fitness of the source subpopulation (**Figures 2** and **S1**) due to long-term differences in population size. Gene flow from the small to the large population (Model 2) only increased the recipient subpopulation’s fitness by 1-2% (*h*=0, **Figure S1**), and thus did not drastically affect the trajectory of *w_R_*/*w_S_* (**Figure 2**). However, gene flow from the large to the small population (Model 3) drastically and immediately increased the fitness of the recipient population (*h*=0, **Figures 2** and **S1**). Because the population size of the recipient subpopulation remained small in Model 3, fitness continued to decline quickly after admixture. In both models, increased fitness in the recipient population after admixture is a consequence of a decrease in the mean number of homozygous deleterious sites per individual due to admixture. These effects are more pronounced for Model 3 because of the higher number of homozygous sites per individual in the large subpopulation (on average about 110; **Figure S3**) compared to the small subpopulation (on average about 50-60; **Figure S3**). In other words, more private recessive deleterious variants are masked by introgressing haplotypes in Model 3, despite the fact that introgressing haplotypes carry a slightly larger number of deleterious variants. Importantly, gene flow had little effect on the mean number of deleterious variants per haplotype (**Figure S2**). Again, our simulations show that the fitness changes from admixture should be largest when recombination rates are low.

Due to these differences in fitness between the small and large subpopulations, the frequency of introgression-derived ancestry in the recipient population changed noticeably for both the additive and recessive models (Models 2 and 3 in **Figure 3**). When fitness effects were additive, these changes were directly linked to selection on introgressed variation. If introgressed haplotypes carried a larger deleterious burden (i.e. came from the smaller population as in Model 2), introgressed ancestry linked to deleterious alleles was removed by selection (long-term *p_I_*≈0-4%). On the other hand, if introgressed haplotypes had a smaller deleterious burden (i.e. came from the larger population as in Model 3), linked introgression-derived ancestry increased in frequency due to selection for haplotypes with fewer deleterious variants (long-term *p_I_*≈6-38%). Again, we observe that the magnitude of this effect is inversely related to the recombination rate. Specifically, the proportion of introgression-derived ancestry decreased or increased at the greatest magnitude in simulations with low recombination, and the least in simulations with a high recombination rate (Models 2 and 3 in **Figure 3**).

When fitness effects were recessive, the frequency of introgression-derived ancestry in the recipient subpopulation were determined by heterosis and differences in genetic load between subpopulations. Gene flow from the large to the small subpopulation (Model 3) resulted in slight (*p_I_*=7%, *r*=10^−6^) to drastic increases (*p_I_*=51%, *r*=10^−9^) in the average proportion of introgression-derived ancestry (Model 3, **Figure 3**). However, admixture from the small to the large (Model 2) population resulted in smaller (*p_I_*≈6-17%) proportions of introgression-derived ancestry (Model 2, **Figure 3**). Additionally, the increase in frequency from selection on recessive variation opposes the effect of selection on additive variation. In the case of Model 3, heterosis and differences in load drive the frequency of introgression-derived ancestry in the same direction, but in the case of Model 2, these factors work in opposite directions. The rate of change of the proportion of introgression-derived ancestry was greatest in simulations with low recombination rates (*r*=10^−9^).

### Long-term population contractions with subsequent population recovery greatly influence the dynamics of introgression

To investigate what occurs when differences in genetic load exist between two subpopulations, but where the strength of selection is increased in the recipient population post-admixture, we simulated another split model where the recipient subpopulation is subjected to a long-term bottleneck but recovers to its original size (Model 4, **Figure 1**). Specifically, the model we simulated was a split model where the recipient subpopulation is reduced to 1,000 diploids immediately after the population split and for 20,000 generations afterwards. At that time point, gene flow occurred in a single generation at an admixture proportion of 5%. Immediately following the admixture event, the recipient subpopulation was restored to its original size (10,000 diploids).

Like the model where the recipient subpopulation size remained small after the split (Model 3), immediately prior to admixture, the recipient population’s fitness was approximately 7-10% less than the fitness of the source population when mutations were additive and 10-30% less than the fitness of the source subpopulation when mutations were recessive (**Figures 2** and **S1**). When mutations were additive, admixture and population recovery result in a gradual increase in the recipient subpopulation’s fitness (*h*=0.5, **Figures 2** and **S1**). When mutations were recessive, admixture rapidly increases the fitness of the recipient subpopulation to at least 90% and up to 95% of its pre-bottleneck fitness, then continues to recover slowly thereafter (*h*=0.0, **Figures 2** and **S1**). In both cases, gene flow results in substantial decreases in the number of deleterious variants per haplotype (**Figure S2**) as well as the number of homozygous deleterious sites per individual (**Figure S3**). The quickest increase in fitness is again observed in the simulations with the lowest recombination rate (*r*=10^−9^; **Figures 2** and **S1**).

Large changes in subpopulation fitness are tied to the largest changes in the frequency of introgression-derived ancestry in the recipient subpopulation (Model 4 in **Figure 3**). When mutations are additive, the fraction of introgression-derived ancestry quickly increases from an initial *p_I_*=5% to *p_I_*≈7-68%, with higher long-term fractions of introgression-derived ancestry occurring at lower recombination rates. When mutations are recessive, the fraction of introgression-derived ancestry has increases to *p_I_*≈10-74%. Thus, the population expansion after the bottleneck drives a rapid increase in the frequency of introgressed ancestry in the recipient population.

### Increasing population split times enhances the effect of heterosis

In models where fitness effects are recessive, heterosis, although modulated by subpopulation differences in genetic load, determines the fitness effects of admixture and the direction of selection on introgression-derived ancestry. However, in these simulations, we have fixed the split time before admixture at 2*N* generations, a substantially long time for private deleterious variation to accumulate within each subpopulation. To further examine the relationship between split time and selection on introgression-derived ancestry, we simulated split models while varying the split time before gene flow. Specifically, we simulated gene flow between two populations of equal size; one where the recipient population experienced a brief bottleneck of 1,000 diploids for the 50 generations immediately before the admixture event; and one where the recipient population’s size was reduced to 1,000 diploids immediately after the split until a single pulse of admixture at 5%, where it then recovered to the original size of 10,000 diploids. These models are analogous to Models 0 and 4 (**Figure 1**), respectively, the only difference being the split time before admixture. The recombination rate was set to *r*=10^−9^ in these simulations.

**Figure 4** depicts the long-term proportion of introgressed ancestry, *p_I_*, 10,000 generations after the single pulse of admixture for these three models varying the amount of time between the split and admixture (*t_s_*). First, we found that across our range of simulated *t_s_, p_I_* always increases monotonically with *t_s_* regardless of the underlying demography (**Figure 4**). This can be attributed to the fact that longer split times result in more deleterious variation being unique to each population (**Figure S4**), enhancing heterosis after admixture. Second, we found as a bottleneck increases in duration, differences in genetic load become a significant contributor to long-term *p_I_*. At a split time and thus bottleneck time of 20,000 generations, the effects of heterosis and the bottleneck increase long-term *p_I_* nearly 2-fold relative to heterosis alone (compare Model 0 to Model 4 in **Figure 4**).

**Figure 4.**
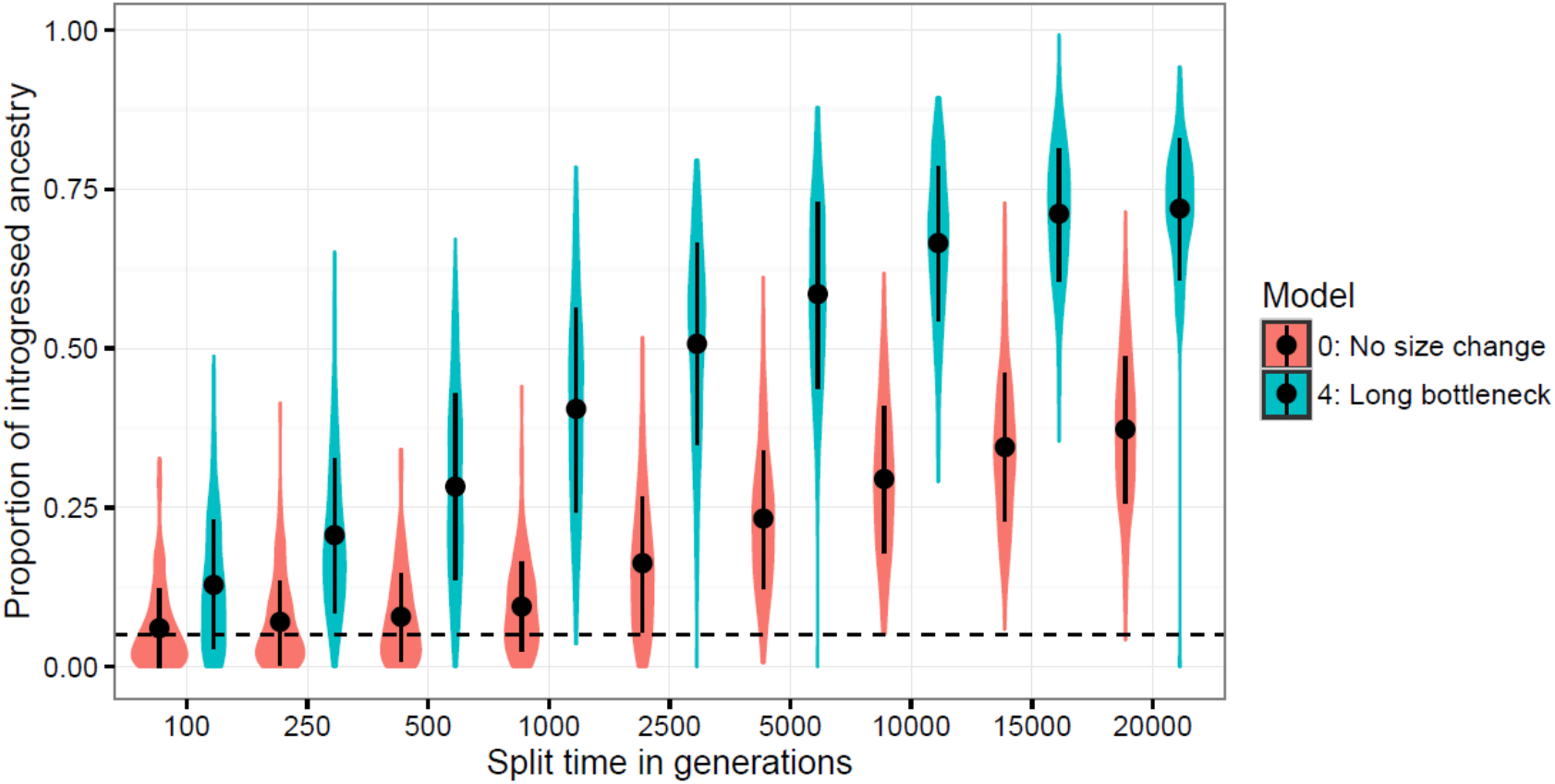
Population split time and population size impact the amount of introgressed ancestry when mutations are recessive. The proportion of ancestry that is introgression-derived, *p_I_*, is shown for 200 simulation replicates and two demographic models (Model 0 and Model 4, refer to **Figure 1**). The recombination rate in all simulations is *r*=10^−9^ per base pair. Violin plots represent the density while dot and whiskers represent the mean and one standard deviation to either side. The horizontal black line represents the initial admixture proportion of 0.05. Note that as the split time increases, *p_I_* also increases.

### A realistic map of chromosomal structure and recombination rates

Thus far we have shown how selection on deleterious variation can affect the dynamics of introgression-derived ancestry when the recombination rate was set to a single value in each set of our simulations. The correlation between introgression-derived ancestry and genomic features such as local recombination rates or exon density are often considered potential evidence for selection against introgression-derived ancestry due to genomic incompatibility or maladaptive alleles (Brandvain et al. 2014; Sankararaman et al. 2014; Pool 2015; Aeschbacher et al. 2017; Corbett-Detig and Nielsen 2017) or adaptive introgression (Corbett-Detig and Nielsen 2017). Selection on deleterious variation is one possible confounder of these patterns (Harris and Nielsen 2016; Juric et al. 2016; Wang et al. 2017).

To investigate how the correlation of introgression-derived ancestry with genomic features is influenced by deleterious variation under different demographic models, we simulated a 100 Mb segment of human chromosome 1, using realistic exon definitions and a recombination map defined on a 10 kb scale (see **Methods**). Unlike the simulations we described previously, we fixed the exon definitions and recombination map to be the same in all simulations. We simulated under three of the split models described previously (**Figure 1**): Model 0, Model 2, and Model 4, separately for recessive and additive fitness effects. Only deleterious mutations were simulated. At the end of each simulation, we split the chromosome into non-overlapping 100 kb windows and computed the frequency of introgression-derived ancestry, exon density, and the average recombination rate in each window.

The average genomic landscape of introgression for 100 simulation replicates varied widely across demographic models and dominance coefficients (**Figure 5** or see **Figure S5** for a representative single simulation replicate). In general, simulations with recessive mutations always showed a genome-wide increase in the frequency of introgressed ancestry, and simulations with additive fitness were dependent upon the demographic model. In the model with equal population sizes (Model 0), we observed no average change in the frequency of introgression-derived ancestry when mutations were additive, but when mutations were recessive we observed a large overall genome-wide increase in the frequency of introgression-derived ancestry (**Figure 5**), with several regions that are at high frequency or fixed in a single simulation replicate (**Figure S5**). Importantly, this increase in frequency is only due to selection on recessive mutations and local variation in recombination rate, since no positively selected mutations were simulated. In the model where introgressed haplotypes contained a larger deleterious burden (Model 2), we observed an overall depletion of introgressed ancestry consistent with the effects of purifying selection upon introgressed ancestry (**Figure 5**). However, for the model with recessive mutations, the effects of heterosis were strong enough such that many genomic regions showed average increases in frequency of 5-10% in our simulations. Importantly, Harris and Nielsen (2016) predicted that heterosis would increase the frequency of introgressed ancestry by only a few percent, but our simulations with a similar demographic model show that low recombination rates can greatly enhance the increase in frequency of introgressed ancestry. Finally, when we simulated the introgression of haplotypes from a population with lower genetic load (Models 3 and 4), we observed drastic, genome-wide increases in the average frequency of introgressed ancestry in the recipient subpopulations (**Figure 5**) as well as many fixed loci in individual simulations (**Figure S5**), regardless of whether fitness effects of mutations were additive or recessive. For example, local regions of the simulated chromosome showed an increase in introgressed ancestry from an initial frequency of 5% up to 70-80% frequency. This is the type of signature that would be unlikely to be generated under neutral demographic models and could be attributed to adaptive introgression.

**Figure 5.**
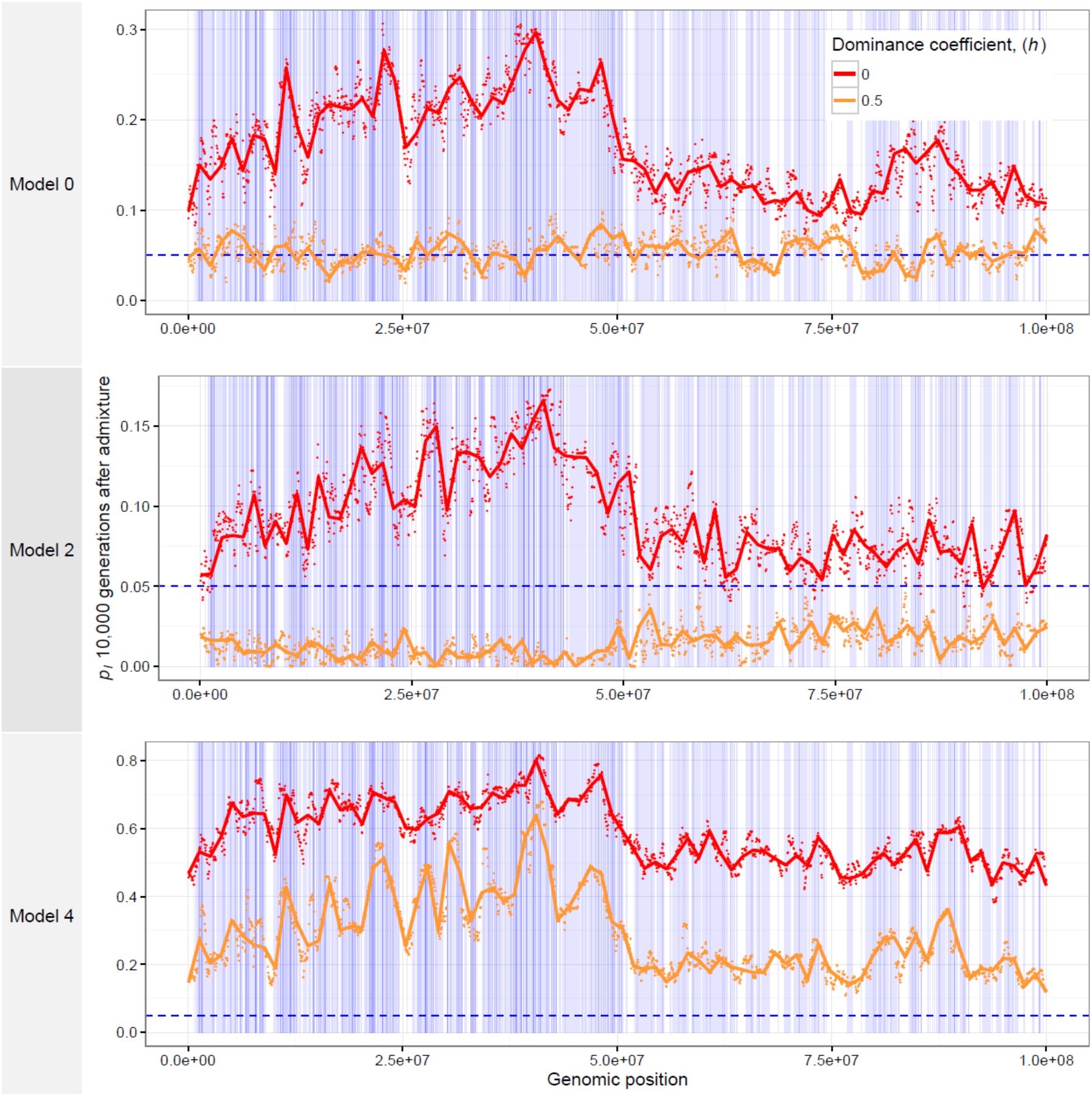
The average genomic landscape of introgression for three demographic models. The frequency of ancestry that is introgression-derived is shown for non-overlapping 100 kb windows in a simulated 100 Mb region of human chromosome 1. The model numbers refer to the models shown in **Figure 1**. Points represent a single value for each 100 kb window and lines are loess curves fitted to the data. The horizontal, blue dashed line represents the initial frequency of introgression-derived ancestry, *p_I_*=0.05. Vertical blue bars represent genes in which deleterious mutations can occur. Red curves denote the results for recessive mutations while orange curves show the results for additive mutations.

It is also notable that the frequency of introgression-derived ancestry (*p_I_*) in each window appears to be driven by exon density, or the local concentration of sites at which deleterious mutations can occur. For recessive mutations, *p_I_* is greatly increased on the left-hand side of the simulated chromosome, which tends to be more gene-rich than the right-hand side of the chromosome (**Figures 5** and **S5**). For additive mutations, the pattern is not as straightforward. When introgressed ancestry is depleted in the recipient population (Model 2), this depletion is strongest in the left-hand side of the figure compared to the right-hand side. In Model 4, where introgressed ancestry increases in the recipient population, the increase is greatest in the most gene-dense part of the chromosome.

We show the correlations that deleterious mutations generate between genomic features and the frequency of introgressed ancestry, measured in 100 kb windows, in **Table 2**. In the model of equal population sizes (Model 0), the frequency of introgression-derived ancestry is not significantly related to the recombination rate or exon density when fitness is additive, but is positively correlated to exon density when fitness effects are recessive. When introgressed ancestry comes from the population with higher load (Model 2), the frequency of introgression-derived ancestry is positively correlated to the recombination rate and negatively correlated to exon density when fitness is additive. When fitness effects are recessive in this model, the frequency of introgressed ancestry is only positively correlated to exon density. Lastly, when introgressed ancestry comes from a larger population having a lower deleterious burden than the recipient population (Model 4), a negative correlation is observed between the frequency of introgression-derived ancestry and recombination rate, and a positive correlation between the frequency of introgression-derived ancestry and exon density. For the last model, these correlations are observed for both models of additive and recessive mutations.

**Table 2.**
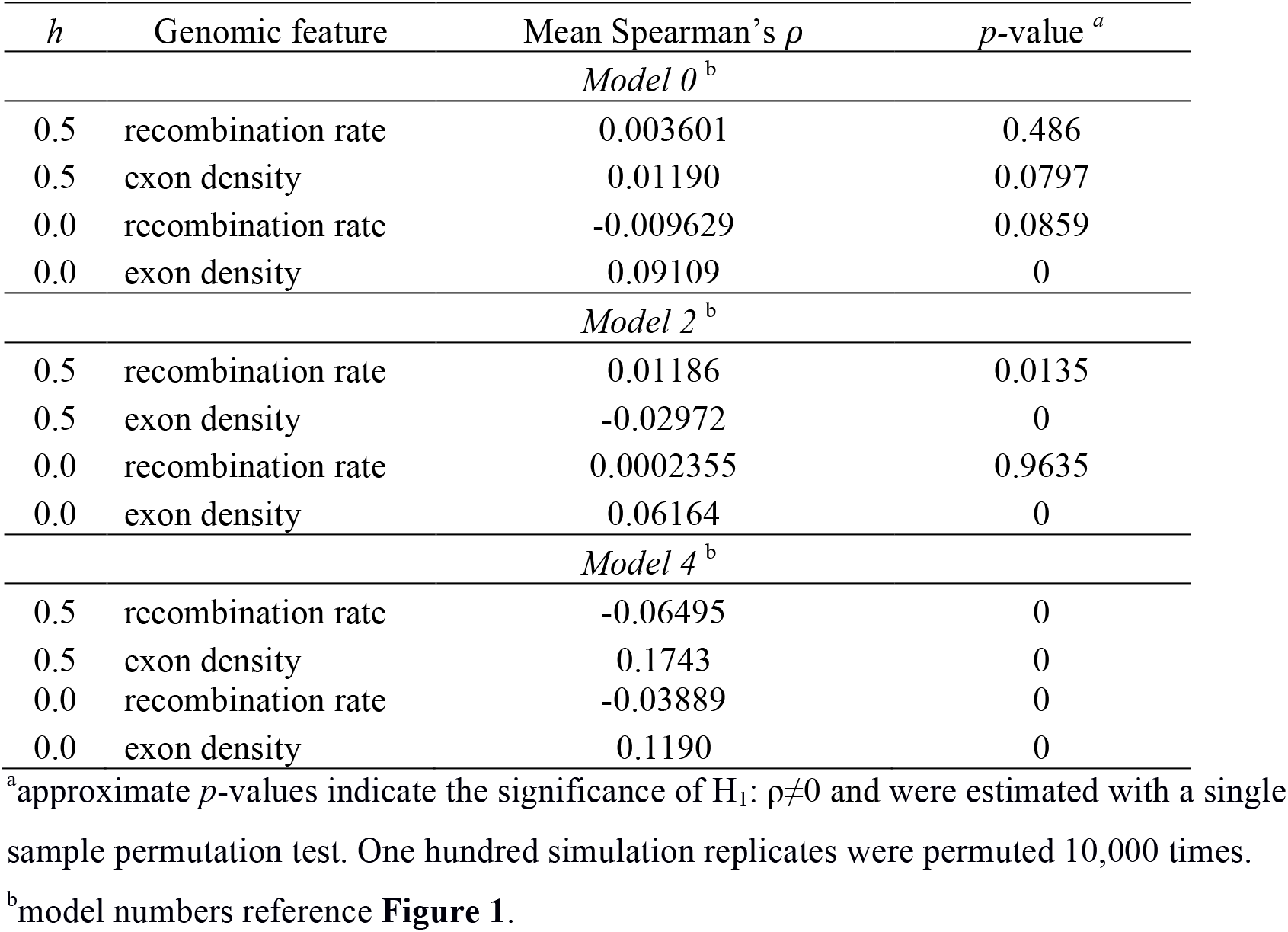
The correlation of recombination rate or exon density with the long-term frequency of introgression-derived ancestry in 100 kb windows is affected by dominance and demography.

## Discussion

We have shown, through simulations, that deleterious variation can greatly influence the dynamics of introgression between admixing populations, in markedly different directions for different demographic scenarios and modes of dominance. In addition, the recombination rate is a key parameter that determines the way in which deleterious variants accumulate between populations, how selection acts on admixed individuals, how selection acts at a locus with admixed ancestry, and ultimately the dynamics of introgression-derived ancestry.

Our work further demonstrates how demography can shape patterns of deleterious variation in different populations. Previous studies have examined the role of population size changes (Kirkpatrick and Jarne 2000; Lohmueller et al. 2008; Simons et al. 2014; Balick et al. 2015; Gravel 2016) and serial founder effect models (Peischl and Excoffier 2015; Henn et al. 2016) on deleterious variation. Interpreting how differences in the distribution of deleterious variation impact fitness has been a contentious issue (Fu et al. 2014; Lohmueller 2014b; Do et al. 2015; Henn et al. 2015; Simons and Sella 2016). In this study, we observed that admixture can increase the fitness of the recipient population, sometimes drastically if the source population is of larger long-term effective size and thus carries lower genetic load. Generally, gene flow is observed to drive smaller, subtle changes in fitness. Nevertheless, the influx of new alleles can result in a rearrangement of deleterious variants in an admixed population (**Figures S2** and **S3**), and even subtle or no changes in fitness or the number of derived alleles per individual can lead to significant shifts in the frequency of introgressed ancestry (e.g. see Model 0, *h*=0, in **Figure 3**). These effects can be long lasting, persisting for thousands of generations in some of our simulations (**Figures 2, 3, S1**). If gene flow or hybridization is a significant feature of a study population, studies concerning load should consider the fitness consequences of admixture as well as population size changes.

That dynamics of introgression-derived ancestry can thus be driven by deleterious variation also is particularly relevant for the study of gene flow between populations or species. Patterns of introgression between hybridizing species are often asymmetric, vary across the genome, and can be driven by demography at expansion fronts (Currat et al. 2008), dispersal processes (Amorim et al. 2017), or by natural selection. However, when natural selection is implicated as driving changes in introgression-derived ancestry, processes such as genomic incompatibility or adaptive introgression are usually invoked to explain variation in introgression across the genome. We have shown that standing deleterious variation, rather than differences in selection on alleles transplanted onto a new genomic background or new environment, has the potential to explain some of these patterns.

To the best of our knowledge, only a few studies have considered the contribution of selection on deleterious variation to observed patterns of introgression (Ingvarsson and Whitlock 2000; Gravel 2016), and mostly in specific systems (Harris and Nielsen 2016; Juric et al. 2016; Wang et al. 2017; Schumer M, Xu C, Powell D, Durvasula A, Skov L, Holland C, Sankararaman S, Andolfatto P, Rosenthal G, Przeworski M, unpublished data, https://www.biorxiv.org/content/early/2017/11/01/212407, accessed Nov. 11, 2017). It is possible that deleterious variation may play a similar role in other species, in particular those that have experienced reductions in population size or range expansions associated with human demography. For example, work by Pool (2015) and Corbett-Detig and Nielsen (2017) on mapping the introgression of African ancestry into North American populations of *Drosophila melanogaster* show that the frequency of introgressed African ancestry is negatively correlated with the recombination rate. This is particularly notable because the opposite relationship is observed for Neanderthal ancestry in humans (Sankararaman et al. 2014) and in hybridizing populations of swordtail fish (Schumer M, Xu C, Powell D, Durvasula A, Skov L, Holland C, Sankararaman S, Andolfatto P, Rosenthal G, Przeworski M, unpublished data, https://www.biorxiv.org/content/early/2017/11/01/212407, accessed Nov. 11, 2017), where regions of the genome with high recombination rate are enriched for introgressed ancestry. Our simulations show that selection on deleterious variants can plausibly explain these opposing patterns in different species.

Originating in Africa, *D. melanogaster* has serially colonized the world in association with humans (Stephan and Li 2006; Duchen et al. 2013), including parts of North America approximately a hundred years ago (Keller 2007). If derived populations of *D. melanogaster* have accumulated differences in deleterious variation due to bottlenecks or increased drift at the expansion front (Peischl and Excoffier 2015; Henn et al. 2016), selection in North American populations may favor introgressed African ancestry simply because this ancestry came from a population with a larger long-term effective size, thus carrying fewer deleterious variants. If recessive deleterious variation also creates heterosis in admixed individuals of North American populations of *D. melanogaster*, the effects of heterosis and population size will be synergistic, further enhancing introgression in genomic regions of low recombination.

Importantly, we do not claim that selection on standing deleterious variation explains all the patterns of introgression in *D. melanogaster* or any other species, but rather that it is a plausible alternative explanation that is important to consider when testing hypotheses about the nature of selection on gene flow. In swordtail fish, recombination rates are positively correlated with the frequency of introgressed ancestry even when the source population has a smaller effective population size than the recipient population (Schumer M, Xu C, Powell D, Durvasula A, Skov L, Holland C, Sankararaman S, Andolfatto P, Rosenthal G, Przeworski M, unpublished data, https://www.biorxiv.org/content/early/2017/11/01/212407, accessed Nov. 11, 2017). This pattern is not explained by models that only include deleterious variation.

Because selection on additive and recessive variation can act in complementary or opposing directions, our study also highlights the fundamental importance of understanding the distribution of selection coefficients and the relationship to dominance coefficients in natural populations (the *h-s* relationship). In this study, we simulated additive and recessive mutations separately, using the same distribution of fitness effects, so that we could demonstrate the effect of only changing the dominance coefficient. However, strongly deleterious new mutations are more likely to be partially or fully recessive (Simmons and Crow 1977; Agrawal and Whitlock 2011; Huber CD, Durvasula A, Hancock AM, Lohmueller KE, unpublished data, https://www.biorxiv.org/content/early/2017/08/31/182865, last accessed Nov. 13, 2017) and so new mutations are likely to have varying degrees of recessivity.

The underlying demographic model will determine how these additive and recessive new mutations should interact after gene flow. For example, the introgression of deleterious haplotypes should be assisted by recessive deleterious mutations but impeded by additive ones, leading to uncertainty about the overall contribution of the effects of deleterious variation in certain scenarios, such as Neanderthal to human admixture (Harris and Nielsen 2016). In other scenarios, selection on additive and recessive variants should operate in the same direction. There is evidence for this type of introgression of wild teosinte into maize (Wang et al. 2017), but it is difficult to disentangle the effects of selection on additive versus recessive variation.

Our simulations reveal that the recombination rate also influences the dynamics of introgressed ancestry in the presence of deleterious variation. Models of Hill-Robertson interference (Hill and Robertson 1966; Keightley and Otto 2006) predict that deleterious mutations will not be removed as effectively in regions of the genome with low recombination rates because they may be linked to the non-deleterious alleles at other sites. We observed the opposite effect in our simulated admixed populations. Specifically, in our simulations, the fitness in the admixed population increased the most for the lowest recombination rates, suggesting that deleterious mutations were most effectively eliminated when recombination rates were the lowest. To understand this effect, it is important to realize that selection for a haplotype will be most effective when all alleles on a haplotype have fitness effects in the same direction. For example, if introgression-derived ancestry carries fewer deleterious variants than the other haplotypes in the recipient population, selection will act to increase the frequency of the protective alleles contained within the introgressed ancestry. This applies directly to our simulations of admixture since immediately following an admixture event, all the protective or deleterious variants are found on the same haplotype. Higher rates of recombination will result in the selected variants being shuffled onto different haplotypes, decreasing the efficacy of selection.

An important objective of genomic studies of hybridization is to identify loci that are adaptively introgressed and to ascertain the overall importance of introgression to adaptive evolution (Racimo et al. 2015). Genomic regions that contain introgressed haplotypes at high frequency are considered likely candidates for adaptive introgression (Huerta-Sánchez et al. 2014; Gittelman et al. 2016; Racimo et al. 2017; Richards and Martin 2017), but we have shown that deleterious variation can generate similar patterns, even in the absence of new beneficial mutations and local adaptation. Model-based statistical approaches that compare summary statistics computed for a particular window of the genome to a simulated null distribution that only accounts for demography may thus be misled by deleterious variation. Again, it may be difficult to differentiate heterosis due to the masking of deleterious recessive alleles from heterozygote advantage at introgressed loci, despite the fact that these are two very different evolutionary processes with dramatically different biological interpretations.

Our results argue that new null models are needed in studies seeking to identify candidates of adaptive introgression. These new null models should include deleterious genetic variation, as well as complex demography. In order for these models to accurately capture the dynamics of deleterious variation, they should also include realistic parameters for the DFE of deleterious mutations and the relationship between dominance coefficients and selection coefficients. Lastly, the new null models should also include realistic models of the variation in recombination rate across the genome, as recombination rate is a key determinant of the dynamics of introgression (**Figure 3**). Failure to consider deleterious variation in a realistic way in studies of admixing populations or hybridizing species can mislead inferences about evolutionary processes acting on the genome.

## Materials and Methods

### Simulation details

All simulations were performed with SLiM 4.2.2 (Haller and Messer 2017). The sequences from simulations with randomly generated chromosome structure were approximately 5 Mb in length (see below). The simulated sequences generated using features from human chromosome 1 were fixed to be exactly 100 Mb in length. Every simulation contained intergenic, intronic, and exonic regions, but only nonsynonymous new mutations experienced natural selection. The per base pair mutation rate was constant and set to *μ*=1.5 × 10^−8^. Within coding sequences, we set nonsynonymous and synonymous mutations to occur at a ratio of 2.31:1 (Huber et al. 2017). The selection coefficients (*s*) of new nonsynonymous mutations were drawn from a gamma-distributed DFE with shape parameter 0.186 and expected selection coefficient E[*s*] = - 0.01314833 (Kim et al. 2017). We chose to simulate additive (*h*=0.5) and recessive (h=0) fitness separately, using the same DFE for *s* for each simulation, to allow the effects of dominance to be directly compared.

Importantly, we chose to discard from our simulations, and therefore from calculations of fitness, mutations that were fixed in the ancestral or both subpopulations. Although fixed deleterious variants contribute to the overall genetic load of finite populations, they will have no effect on the relative differences between admixing subpopulations and no effect on the dynamics of introgression-derived ancestry. Therefore, each fitness calculation does not reflect the true fitness, but rather the fitness components that are relevant during gene flow.

An admixture event in SLiM is handled by modifying the way the parents in each generation are chosen (SLiM manual 5.2.1). For example, at an admixture proportion of 5% the recipient population reproduces as follows. Five percent of the parents of the recipient population, in that generation, are chosen from the source population, and 95% of the parents are chosen from the recipient population.

### Simulations with randomly generated chromosomal structure

Unless specified otherwise, the chromosomal structure of each simulation was randomly generated by drawing exon lengths from *Lognormal*(*μ* = log(50), *σ*^2^ = log(2)), intron lengths from *Lognormal*(*μ* = log(100), *σ*^2^ = log(1.5)), and the length of noncoding regions from *unif*(100,5000), following the specification in the SLiM 4.2.2 manual (7.3), which is modeled after the distribution of intron and exon lengths in Deutsch and Long (1999). The per-base pair recombination rate (*r*) was fixed in each simulation, but we varied *r* between different sets of simulations where *r*∈{10^−6^,10^−7^,10^−8^,10^−9^}. Lastly, we simulated 200 replicates for each set of simulations, of each specific *h* and *r*.

### Simulations with fixed, realistic chromosomal structure

In simulations with fixed chromosomal structure (**Figure 6**), we fixed the structure to 100 Mb randomly chosen from human genome build GRCh37, chromosome 1 (chr1:5,005,669-105,005,669). The exon ranges were defined by the GENCODE v14 annotations (Harrow et al. 2012) and the sex-averaged recombination map by Kong et al. (2010), averaged over a 10 kb scale.

### Avoiding heterosis in the additive fitness model

Computing fitness as additive at a locus but multiplicative across loci creates artificial heterosis in admixed individuals. This occurs because the product of a fitness decrease will reduce fitness less than the sum of a fitness decrease. As an example, assume two deleterious alleles are in a single individual, each with selection coefficient *s* where *s*<0. If they are found as a single homozygous site, the fitness decrease is usually computed as (1+2*s*). If they are found in two heterozygous sites, the fitness would be computed as (1+*s*)^2^. This second quantity is larger than the first by *s*^2^. Because admixed individuals are more likely to carry deleterious alleles as heterozygotes than non-admixed individuals, the fitness of the admixed individuals will always be higher than a non-admixed individual in the above computation of fitness even when the number of deleterious variants per individual is the same.

Because our intent with an additive fitness model was to make the fitness effect of each variant independent of its genotypic state, we computed additive fitness as purely multiplicative across all deleterious variants, such that an individual *j* carrying *i* variants each with selection coefficient *s_i_* has fitness *w_i_*:

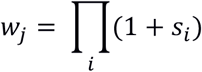

This computation of fitness is approximately equivalent to additive fitness. Recessive fitness effects were computed in the standard manner, that is, as 1+2*s_i_* when homozygous for the deleterious allele and as 1 otherwise.

### Data availability

All scripts necessary for reproducing the simulations we have presented are available at https://github.com/LohmuellerLab/admixture_load_scripts.

## Acknowledgements

We thank Jacqueline Robinson, Annabel Beichman, and other members of the Lohmueller Lab for helpful comments throughout the project. This work was supported by the National Institutes of Health (R35 GM119856 to K.E.L.).

## References

Aeschbacher, S, Selby JP, Willis JH, Coop G. 2017. Population-genomic inference of the strength and timing of selection against gene flow. Proc. Natl. Acad. Sci. 114:7061–7066.

Agrawal AF, Whitlock MC. 2011. Inferences about the distribution of dominance drawn from yeast gene knockout data. Genetics 187:553–566.

Amorim CEG, Hofer T, Ray N, Foll M, Ruiz-Linares A, Excoffier L. 2017. Long-distance dispersal suppresses introgression of local alleles during range expansions. Heredity 118:135–142.

Balick DJ, Do R, Cassa CA, Reich D, Sunyaev SR. 2015. Dominance of Deleterious Alleles Controls the Response to a Population Bottleneck. PLOS Genet. 11:e1005436.

Brandvain, Y, Kenney AM, Flagel L, Coop G, Sweigart AL. 2014. Speciation and Introgression between *Mimulus nasutus* and *Mimulus guttatus.* PLOS Genet. 10:e1004410.

Brandvain, Y, Wright SI. 2016. The Limits of Natural Selection in a Nonequilibrium World. Trends Genet. 32:201–210.

Casals, F, Hodgkinson A, Hussin J, Idaghdour Y, Bruat V, Maillard T de, Grenier J-C, Gbeha E, Hamdan FF, Girard S, et al. 2013. Whole-Exome Sequencing Reveals a Rapid Change in the Frequency of Rare Functional Variants in a Founding Population of Humans. PLOS Genet. 9:e1003815.

Corbett-Detig R, Nielsen R. 2017. A Hidden Markov Model Approach for Simultaneously Estimating Local Ancestry and Admixture Time Using Next Generation Sequence Data in Samples of Arbitrary Ploidy. PLOS Genet. 13:e1006529.

Crow JF. 1948. Alternative Hypotheses of Hybrid Vigor. Genetics 33:477–487.

Currat M, Ruedi M, Petit RJ, Excoffier L. 2008. The hidden side of invasions: massive introgression by local genes. Evol. Int. J. Org. Evol. 62:1908–1920.

Deutsch M, Long M. 1999. Intron-exon structures of eukaryotic model organisms. Nucleic Acids Res. 27:3219–3228.

Do R, Balick D, Li H, Adzhubei I, Sunyaev S, Reich D. 2015. No evidence that selection has been less effective at removing deleterious mutations in Europeans than in Africans. Nat. Genet. 47:126–131.

Duchen P, Zivkovic D, Hutter S, Stephan W, Laurent S. 2013. Demographic Inference Reveals African and European Admixture in the North American *Drosophila melanogaster* Population. Genetics 193:291–301.

Fu W, Gittelman RM, Bamshad MJ, Akey JM. 2014. Characteristics of Neutral and Deleterious Protein-Coding Variation among Individuals and Populations. Am. J. Hum. Genet. 95:421–436.

Gazave E, Chang D, Clark AG, Keinan A. 2013. Population Growth Inflates the Per-Individual Number of Deleterious Mutations and Reduces Their Mean Effect. Genetics 195:969–978.

Gittelman RM, Schraiber JG, Vernot B, Mikacenic C, Wurfel MM, Akey JM. 2016. Archaic Hominin Admixture Facilitated Adaptation to Out-of-Africa Environments. Curr. Biol. 26:3375–3382.

Gravel S. 2016. When Is Selection Effective? Genetics 203:451–462.

Haller BC, Messer PW. 2017. SLiM 2: Flexible, Interactive Forward Genetic Simulations. Mol. Biol. Evol. 34:230–240.

Harris K, Nielsen R. 2016. The Genetic Cost of Neanderthal Introgression. Genetics 203:881–891.

Harrow J, Frankish A, Gonzalez JM, Tapanari E, Diekhans M, Kokocinski F, Aken BL, Barrell D, Zadissa A, Searle S, et al. 2012. GENCODE: The reference human genome annotation for The ENCODE Project. Genome Res. 22:1760–1774.

Henn BM, Botigué LR, Bustamante CD, Clark AG, Gravel S. 2015. Estimating the mutation load in human genomes. Nat. Rev. Genet. 16:333–343.

Henn BM, Botigué LR, Peischl S, Dupanloup I, Lipatov M, Maples BK, Martin AR, Musharoff S, Cann H, Snyder MP, et al. 2016. Distance from sub-Saharan Africa predicts mutational load in diverse human genomes. Proc. Natl. Acad. Sci. 113:E440–E449.

Hill WG, Robertson A. 1966. The effect of linkage on limits to artificial selection. Genet. Res. 8:269–294.

Huber CD, Kim BY, Marsden CD, Lohmueller KE. 2017. Determining the factors driving selective effects of new nonsynonymous mutations. Proc. Natl. Acad. Sci. 114:4465–4470.

Huerta-Sánchez E,Jin X, Asan, Bianba Z, Peter BM, Vinckenbosch N, Liang Y, Yi X, He M, Somel M, et al. 2014. Altitude adaptation in Tibetans caused by introgression of Denisovan-like DNA. Nature 512:194–197.

Ingvarsson PK, Whitlock MC. 2000. Heterosis increases the effective migration rate. Proc. R. Soc. Lond. B Biol. Sci. 267:1321–1326.

Juric I, Aeschbacher S, Coop G. 2016. The Strength of Selection against Neanderthal Introgression. PLOS Genet. 12:e1006340.

Keightley PD, Otto SP. 2006. Interference among deleterious mutations favours sex and recombination in finite populations. Nature 443:89–92.

Keller A. 2007. Drosophila melanogaster’s history as a human commensal. Curr. Biol. 17:R77–R81.

Kim BY, Huber CD, Lohmueller KE. 2017. Inference of the Distribution of Selection Coefficients for New Nonsynonymous Mutations Using Large Samples. Genetics 206:345–361.

Kimura M, Maruyama T, Crow JF. 1963. The Mutation Load in Small Populations. Genetics 48:1303–1312.

Kirkpatrick M, Jarne P. 2000. The Effects of a Bottleneck on Inbreeding Depression and the Genetic Load. Am. Nat. 155:154–167.

Kong A, Thorleifsson G, Gudbjartsson DF, Masson G, Sigurdsson A, Jonasdottir A, Walters GB, Jonasdottir A, Gylfason A, Kristinsson KT, et al. 2010. Fine-scale recombination rate differences between sexes, populations and individuals. Nature 467:1099–1103.

Liu Q, Zhou Y, Morrell PL, Gaut BS. 2017. Deleterious Variants in Asian Rice and the Potential Cost of Domestication. Mol. Biol. Evol. 34:908–924.

Lohmueller KE. 2014a. The Impact of Population Demography and Selection on the Genetic Architecture of Complex Traits. PLOS Genet 10:e1004379.

Lohmueller KE. 2014b. The distribution of deleterious genetic variation in human populations. Curr. Opin. Genet. Dev. 29:139–146.

Lohmueller KE, Indap AR, Schmidt S, Boyko AR, Hernandez RD, Hubisz MJ, Sninsky JJ, White TJ, Sunyaev SR, Nielsen R, et al. 2008. Proportionally more deleterious genetic variation in European than in African populations. Nature 451:994–997.

Marsden CD, Vecchyo DO-D, O’Brien DP, Taylor JF, Ramirez O, Vilà C, Marques-Bonet T, Schnabel RD, Wayne RK, Lohmueller KE. 2016. Bottlenecks and selective sweeps during domestication have increased deleterious genetic variation in dogs. Proc. Natl. Acad. Sci. 113:152–157.

Moyers BT, Morrell PL, McKay JK. 2017. Genetic costs of domestication and improvement. J. Hered. esx069, https://doi.org/10.1093/jhered/esx069

Muller HJ. 1964. The relation of recombination to mutational advance. Mutat. Res. Mol. Mech. Mutagen. 1:2–9.

Payseur BA, Rieseberg LH. 2016. A genomic perspective on hybridization and speciation. Mol. Ecol. 25:2337–2360.

Pedersen C-ET, Lohmueller KE, Grarup N, Bjerregaard P, Hansen T, Siegismund HR, Moltke I, Albrechtsen A. 2017. The Effect of an Extreme and Prolonged Population Bottleneck on Patterns of Deleterious Variation: Insights from the Greenlandic Inuit. Genetics 205:787–801.

Peischl S, Excoffier L. 2015. Expansion load: recessive mutations and the role of standing genetic variation. Mol. Ecol. 24:2084–2094.

Pool JE. 2015. The Mosaic Ancestry of the Drosophila Genetic Reference Panel and the D. melanogaster Reference Genome Reveals a Network of Epistatic Fitness Interactions. Mol. Biol. Evol. 32:3236–3251.

Racimo F, Marnetto D, Huerta-Sánchez E. 2017. Signatures of Archaic Adaptive Introgression in Present-Day Human Populations. Mol. Biol. Evol. 34:296–317.

Racimo F, Sankararaman S, Nielsen R, Huerta-Sánchez E. 2015. Evidence for archaic adaptive introgression in humans. Nat. Rev. Genet. 16:359–371.

Richards EJ, Martin CH. 2017. Adaptive introgression from distant Caribbean islands contributed to the diversification of a microendemic adaptive radiation of trophic specialist pupfishes. PLOS Genet. 13:e1006919.

Sankararaman S, Mallick S, Dannemann M, Prüfer K, Kelso J, Pääbo S, Patterson N, Reich D. 2014. The genomic landscape of Neanderthal ancestry in present-day humans. Nature 507:354–357.

Simmons MJ, Crow JF. 1977. Mutations affecting fitness in Drosophila populations. Annu. Rev. Genet. 11:49–78.

Simons YB, Sella G. 2016. The impact of recent population history on the deleterious mutation load in humans and close evolutionary relatives. Curr. Opin. Genet. Dev. 41:150–158.

Simons YB, Turchin MC, Pritchard JK, Sella G. 2014. The deleterious mutation load is insensitive to recent population history. Nat. Genet. 46:220–224.

Stephan W, Li H. 2006. The recent demographic and adaptive history of *Drosophila melanogaster.* Heredity 98:65–68.

Vernot B, Tucci S, Kelso J, Schraiber JG, Wolf AB, Gittelman RM, Dannemann M, Grote S, McCoy RC, Norton H, et al. 2016. Excavating Neandertal and Denisovan DNA from the genomes of Melanesian individuals. Science:aad9416.

Wall JD, Yoshihara Caldeira Brandt D. 2016. Archaic admixture in human history. Curr. Opin. Genet. Dev. 41:93–97.

Whitlock MC, Ingvarsson PK, Hatfield T. 2000. Local drift load and the heterosis of interconnected populations. Heredity 84 (Pt 4):452–457.

